# Ena/VASP proteins mediate endothelial cell repulsion from ephrin ligands

**DOI:** 10.1101/2023.04.19.537567

**Authors:** Joana Zink, Timo Frömel, Ilka Wittig, Ingrid Fleming, Peter M. Benz

## Abstract

The interaction of Eph receptor tyrosine kinases with their transmembrane ligands; the ephrins, is important for the regulation of cell-cell communication. Ephrin-Eph signaling is probably best known for the discrimination of arterial and venous territories by repulsion of venous endothelial cells away from those with an arterial fate. Ultimately, cell repulsion is mediated by initiating the collapse of the actin cytoskeleton in membrane protrusions. Here, we investigated the role of the Ena/VASP family of actin binding proteins in endothelial cell repulsion initiated by ephrin ligands. Human endothelial cells dynamically extended sheet-like lamellipodia over ephrin-B2 coated surfaces. While lamellipodia of control siRNA transfected cells rapidly collapsed, resulting in a pronounced cell repulsion from the ephrin-B2 surfaces, the knockdown of Ena/VASP proteins impaired the cytoskeletal collapse of membrane protrusions and the cells no longer avoided the repulsive surfaces. Mechanistically, ephrin-B2 stimulation elicited the EphB-mediated tyrosine phosphorylation of VASP, which abrogated its interaction with the focal adhesion protein Zyxin. Nck2 was identified as a novel VASP binding protein, which only interacted with the tyrosine phosphorylated VASP protein. Nck links Eph-receptors to the actin cytoskeleton. Therefore, we hypothesize that Nck-Ena/VASP complex formation is required for actin reorganization and/or Eph receptor internalization downstream of ephrin-Eph interaction in endothelial cells, with implications for endothelial navigation and pathfinding.

## Introduction

An important molecular mechanism that mediates short-distance cell–cell communication is the interaction of the Eph receptor tyrosine kinases with their transmembrane ephrin ligands. In the classical model of ‘forward’ signaling, ephrins act as *in trans* ligands of Eph receptors expressed by neighboring cells, to induce signaling pathways that trigger cell repulsion. In the ‘reverse’ signaling mode, Eph receptors serve as *in trans* as ligands for ephrins, which can elicit both cell repulsion or adhesion (Kania and Klein, 2016). This Ephrin–Eph signaling pathway is rapid and regulates many processes that involve fast changes in cellular motility and/or morphology. While originally discovered during embryonic development in the patterning of the nervous system and axon guidance, ephrin-Eph signaling is also an essential regulator of many vascular processes (Kania and Klein, 2016; Sawamiphak et al., 2010; Vreeken et al., 2020). Most prominent is the interaction of ephrin-B2 with its receptor; EphB4, that regulates the development of blood and lymphatic vessels and helps to establish borders between arterial and venous territories by repulsion of venous endothelial cells away from those with an arterial fate (Kania and Klein, 2016). Ephrin-Eph forward signaling ultimately works by triggering cytoskeletal collapse, to stimulate the retraction of actin-rich membrane protrusions i.e., lamellipodia and filopodia, to mediate cell repulsion. However, underlying mechanisms are elusive and proteins involved in cytoskeletal collapse downstream of Eph receptors and their role in endothelial repulsion and pathfinding are currently not well understood.

The enabled/vasodilator-stimulated phosphoprotein (Ena/VASP) proteins are important mediators of cytoskeletal control, linking kinase signaling pathways to actin assembly. The proteins are localized at sites of high actin turnover, including focal adhesions, the leading edge of lamellipodia, and the tips of filopodia, where they promote actin polymerization and cell migration (Faix and Rottner, 2022). In mammals, the Ena/VASP family of proteins consists of mammalian enabled (Mena), VASP and Ena-VASP-like protein (EVL). The family members share a tripartite domain organization of an N-terminal Ena/VASP homology 1 (EVH1) domain, a central proline-rich region (PRR), and an EVH2 domain at the C terminus. The EVH1 domain mediates subcellular targeting of Ena/VASP by binding to focal adhesion proteins such as vinculin and zyxin (Sechi and Wehland, 2004). In endothelial cells, Ena/VASP proteins are required for intercellular adhesion and barrier function and the proteins are also important regulators of angiogenic sprouting (Benz et al., 2008; Benz et al., 2019; Zink et al., 2021). Given their role at the crossroads of kinase signaling pathways and actin cytoskeleton remodeling, it is tempting to speculate, that Ena/VASP proteins may regulate endothelial cell repulsion or pathfinding downstream of ephrin-Eph receptor signaling. So far, however, there is only circumstantial evidence to support this hypothesis. Ena/VASP proteins colocalize with activated EphB receptors and have been implicated in the repulsion of neural crest cells from ephrin ligands (Evans et al., 2007). Moreover, tyrosine phosphorylated (Y16 and Y39) VASP peptides were detected by high throughput mass spectrometry studies of HEK cells with activated ephrin/EphB signaling (Jorgensen et al., 2009). This study set out to investigate the role of Ena/VASP proteins in endothelial cell repulsion from ephrin ligands and to elucidate potential underlying mechanisms.

## Results and discussion

The impact of Ena/VASP proteins in ephrin-Eph signaling was studied *in vitro* in human umbilical vein endothelial cells using an approach originally designed for analysis of axonal guidance (Knoll et al., 2007). As EVL was expressed at low levels in cells studied, experiments focused on Mena and VASP. The latter proteins were readily detectable and it was even possible to detect the two Mena variants described by others (Gertler et al., 1996). Also, a small interfering (si) RNA approach was successfully used to decrease the expression of Mena and VASP (Figure 1A). Next, the Mena/VASP-expressing or - deficient cells were seeded onto slides coated in strips with immobilized clustered ephrin B2 protein or a control (Figure 1A). The integrity of the stripes was routinely confirmed by incorporation of fluorescent Cy3 dye (Figure 1B) before repulsion was studied by time-lapse microscopy (Figure 1 and supplementary movies 1 and 2). When seeded onto the slide, endothelial cells that expressed Mena and VASP actively avoided the substrate-bound ephrin-B2, resulting in the establishment of endothelial cell stripes on the control matrix (Figure 1C, E, G and I). The endothelial cells were highly motile and dynamically extended sheet-like lamellipodia over the ephrin-B2 stripes but these rapidly collapsed on the repulsive surface to form optically dense structures and membrane ruffles (arrows in Figure 1E and supplementary movie 1). Cells underwent repeated cycles of protrusion and retraction of the leading edge to avoid contact with the immobilized ephrin-B2. Endothelial cells that lacked Mena and VASP failed to align along the stripes in this manner and remained highly motile in all directions (Figure 1 D, F, H and J; and supplementary movie 2). Indeed, lamellipodia extension and morphology on ephrin-B2 and control stripes appeared similar (arrow heads in Figure 1F), and cells were found randomly located over the entire slide. These observations suggest an important role of Ena/VASP proteins in endothelial pathfinding downstream of ephrin-B2–EphB4 signaling and our findings corroborate a previous report that showed a significantly decreased repulsion of VASP-deficient and Mena/VASP-double deficient fibroblasts from substrate-bound ephrin-B2 (Evans et al., 2007).

**Figure 1.**
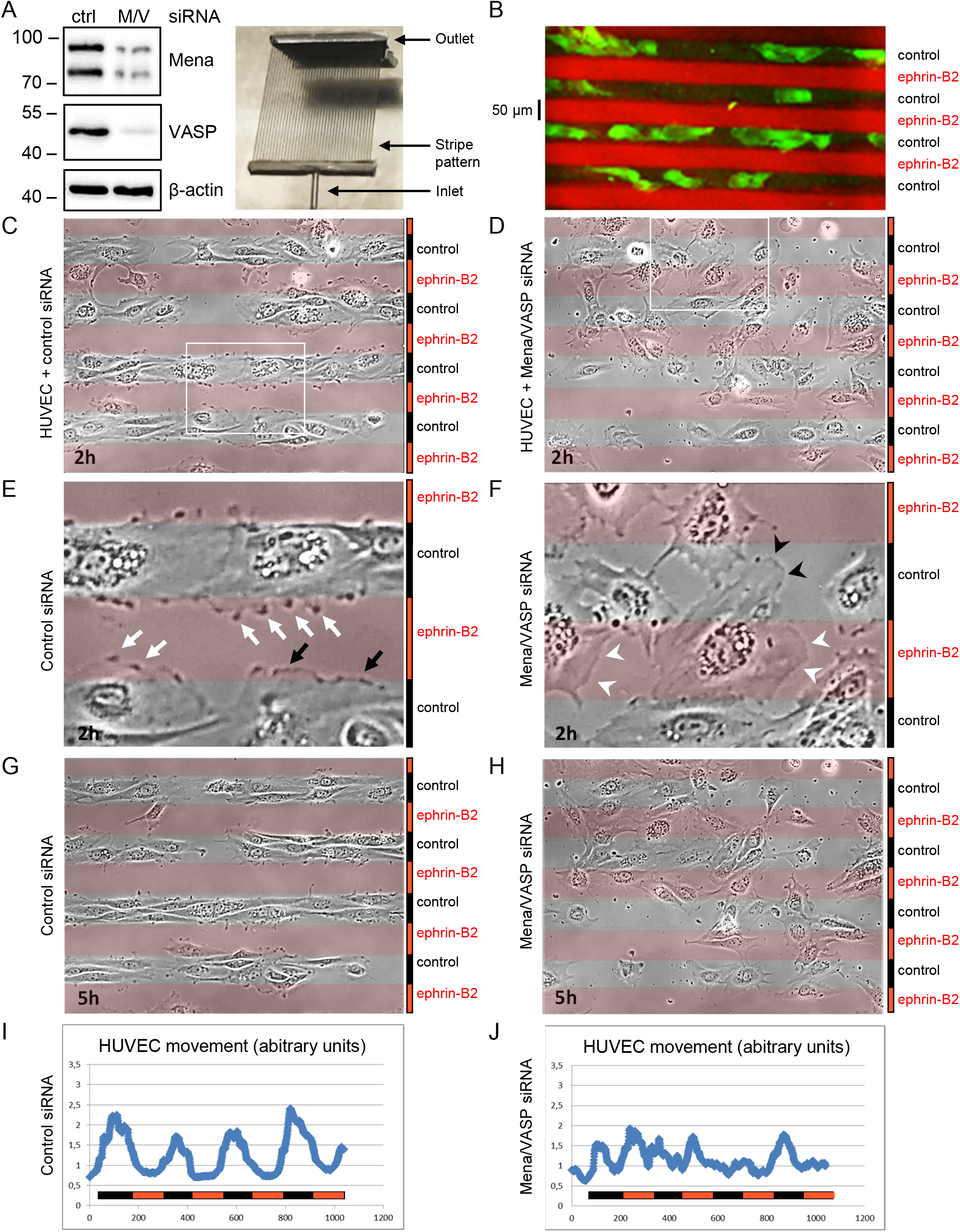
Role of Ena/VASP proteins in endothelial cell repulsion from ephrin ligands. HUVECs, treated with Mena/VASP or control siRNA (**A**), were seeded on alternating 50 μm stripes with clustered ephrin-B2 (labeled with Cy3 fluorescent dye) or control (**B-H**) and imaged by time-lapse microcopy. The image in (A) shows a silicon matrix used to generate the stripes. Representative phase contrast images of cells approximately two hours (**C-F**) and five hours (**G, H**) after seeding are shown; ephrin-B2 stripes are indicated by red overlays. Please note the cytoskeletal collapse in control siRNA-transfected HUVECs on ephrin-B2 stripes, resulting in dot-like structures and membrane ruffles (E, white and black arrows, respectively), whereas no cytoskeletal collapse was observed in Mena/VASP siRNA transfected cells on ephrin-B2 stripes (F, compare lamellipodia indicated by black and white arrowheads, respectively). Cell movement on ephrin-B2 or control stipes in the time-lapse movies was analyzed with Image J (**I, J**). While control siRNA-transfected cells clearly avoided migration on ephrin-B2 coated stripes (I), this repulsive effect was largely abolished in Mena/VASP siRNA-transfected cells (J).

To elucidate the molecular mechanism(s) underlying the Ena/VASP-dependent repulsion of endothelial cells from ephrin-B2 stripes, experiments focused on detecting early changes in EphB signaling. One of the first forward signaling events is the activation of the tyrosine kinase activity of the Eph receptor, which is crucial for Eph-mediated cellular responses, cytoskeletal rearrangements and the regulation of boundary formation (Kania and Klein, 2016). Functionally, serine/threonine phosphorylation events have been shown to control the subcellular targeting of Ena/VASP proteins and their ability to modulate actin dynamics, at least in part by regulating their protein-protein interactions (Barzik et al., 2005; Benz et al., 2008; Benz et al., 2009; Blume et al., 2007). Notably, VASP phosphorylation status is used in numerous studies to assess PKA/PKG-dependent effects in cardiovascular cells and in clinical diagnosis of platelet reactivity (Aleil et al., 2005; Benz and Fleming, 2016; Rukoyatkina et al., 2017; Walter and Gambaryan, 2009). However, little is known about the kinases and functional consequences of Ena/VASP tyrosine phosphorylation. High throughput proteomic studies identified the phosphorylation of Ena/VASP proteins on Y16 and/or Y39 within the EVH1 domains of the proteins ((Helou et al., 2013; Jorgensen et al., 2009; Rikova et al., 2007) and Figure 2A). It was possible to confirm the phosphorylation of VASP Y39 by a ternary complex with c-Abl and Abi-1, and a phosphomimetic Y39D-VASP mutant failed to efficiently localize to focal adhesions (Maruoka et al., 2012). To determine whether VASP tyrosine phosphorylation could be stimulated by ephrins, EphB4-expressing primary human umbilical vein and artery endothelial cells (HUVEC and HUAEC) (Bochenek et al., 2010) as well as EphB4-positive K562 chronic myeloid leukemia cells (Merchant et al., 2017) were treated with pre-clustered ephrin-B2, to elicit strong forward signaling (Kania and Klein, 2016). Immunoprecipitation with VASP-specific antibodies and Western blotting with phosphotyrosine-specific antibodies revealed a pronounced tyrosine phosphorylation of VASP in ephrin-B2 treated cells (Figure 2B), corroborating a previous report that showed colocalization of Ena/VASP proteins with activated Eph receptors in fibroblasts (Evans et al., 2007).

**Figure 2.**
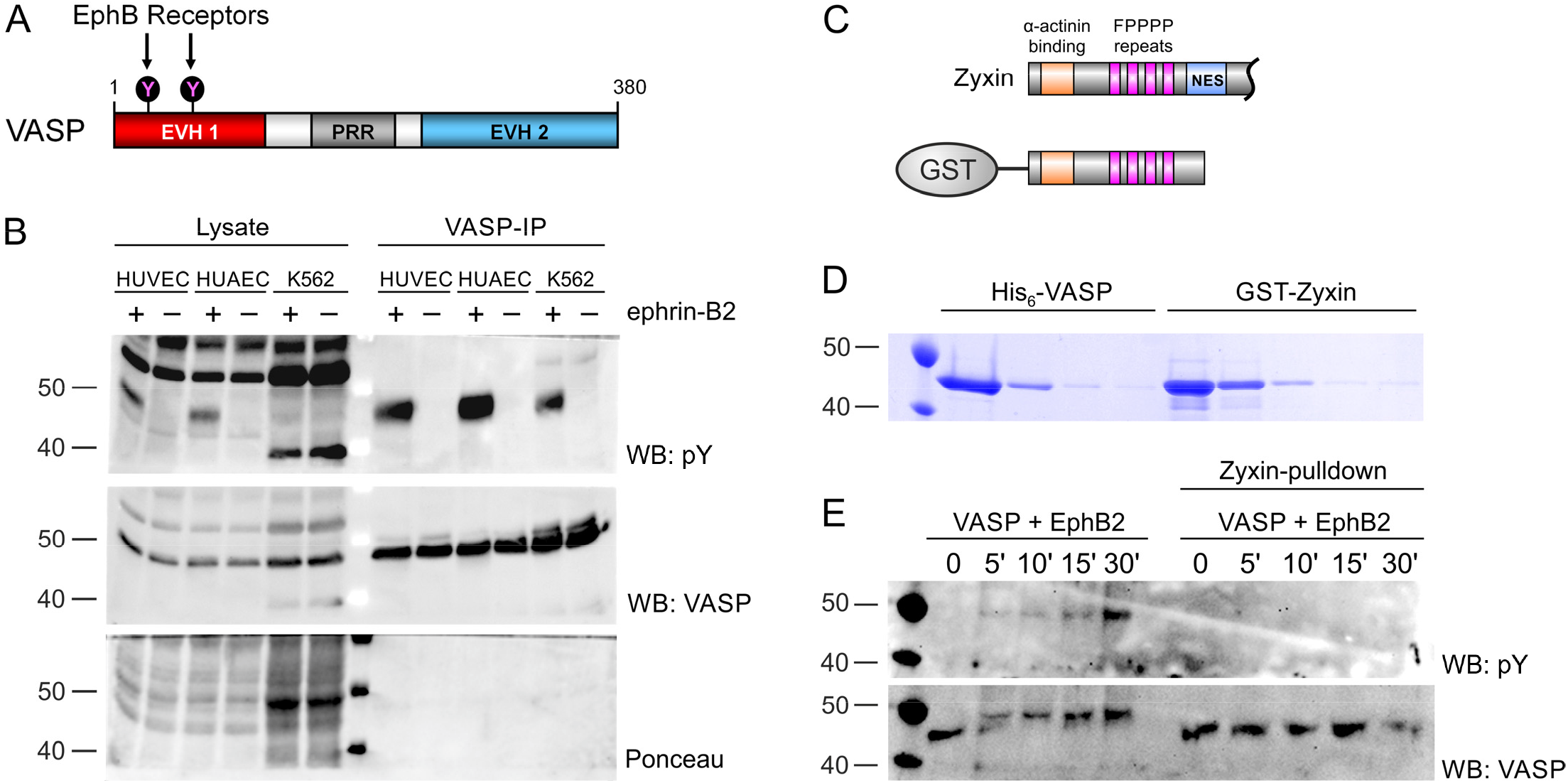
EphB induced VASP tyrosine phosphorylation abrogates zyxin-VASP binding. (**A**) VASP contains two tyrosine residues in the EVH1 domain, which mediates binding to the focal adhesion protein zyxin. (**B**) Stimulation of human umbilical vein or artery endothelial cells (HUVECs, HUAECs), and K562 cancer cells with pre-clustered ephrin-B2 activates EphB receptors and induces VASP tyrosine phosphorylation. VASP was immunoprecipitated and probed with phosphotyrosine-specific antibodies (pY, upper panel). VASP-specific antibodies (middle panel) and ponceau-S staining (lower panel) served as control. (**C**) Scheme and Coomassie stained gel (**D**) of the purified GST-zyxin fusion protein (and VASP) used for the pull-down experiments shown in (E). The ActA repeats (magenta) in zyxin are the binding sites for the VASP EVH1 domain. (**E**) GST pull-down assay with GST-zyxin and in vitro phosphorylated VASP. VASP was in vitro tyrosine phosphorylated or not with recombinant EphB2 for 5-30 minutes. VASP tyrosine phosphorylation was confirmed by Western blotting with phospho-tyrosine specific (pY) monoclonal antibodies, showing a time dependent increase in phosphorylation (upper panel, left). The tyrosine phosphorylated or non-phosphorylated VASP was used in pull-down assays with GST-tagged zyxin. While GST-zyxin readily pull-down VASP (right lower panel), no tyrosine phosphorylated VASP was detected in the precipitated material (upper panel, right), indicating that VASP tyrosine phosphorylation by recombinant EphB2 blocks interaction with the GST-zyxin.

Although we did not assess exactly which tyrosine residue was phosphorylated, we focused on the implication of tyrosine phosphorylation on interaction between the VASP EVH1 domain and the focal adhesion protein zyxin. This was because Y16/39-phosphorylations in the EVH1 domain were picked up in a series of high throughput proteomic screens, including a study with activated ephrin/EphB signaling in HEK cells (Helou et al., 2013; Jorgensen et al., 2009; Rikova et al., 2007). Notably, Y16 is implicated in forming the hydrophobic docking site for EVH1 ligands (Ball et al., 2002; Prehoda et al., 1999). To assess the influence of tyrosine phosphorylation on EVH1 binding to zyxin, pull-down experiments were performed with purified, non-phosphorylated or *in vitro* phosphorylated VASP and a GST-zyxin fusion protein that contained VASP binding sites (Drees et al., 2000) and Figure 2C and D). *In vitro* phosphorylation was performed with recombinant EphB2, which is also a natural receptor for ephrin-B2. This approach revealed that VASP interacted with GST-zyxin in the pull-down assay but only when it was not phosphorylated (Figure 2E). These observations suggest that VASP tyrosine phosphorylation may be a molecular switch that modulates the targeting of Ena/VASP to focal adhesions. This is especially relevant since the impaired recruitment of Ena/VASP proteins to focal adhesions is associated with changes in cell motility (Hoffman et al., 2006).

Nevertheless, reduced focal adhesion targeting of Ena/VASP proteins alone seemed unlikely to explain the results rapid protrusion and retraction of lamellipodia from the stripe assay. Indeed, the latter events are largely independent of the association of lamellipodia to the substratum through (nascent) focal adhesions (Faix and Rottner, 2022). To better understand the implication of Ena/VASP tyrosine phosphorylation we performed phospho-proteomics using tyrosine-phosphorylated or non-phosphorylated VASP as bait. Given that EphB4-mediated VASP phosphorylation is ligand dependent and difficult to control, we used an alternative approach and used two engineered Abl kinases. We generated a hyperactive GKP-Abl kinase mutant, which combines four activating point mutations, as well as a kinase dead (KD) variant (Figure 3A and (Benz et al., 2019)). After co-expression of VASP, Abi-1 and GKP-Abl or KD-Abl in HEK cells, VASP and phospho-tyrosine VASP was purified (Figure 3B) and the phosphorylation status of the proteins was confirmed with phosphotyrosine specific antibodies and by mass spectrometry (Figure 3C-D). After coupling to an affinity matrix, VASP and phospho-tyrosine VASP were used to precipitate VASP binding proteins from endothelial cell lysates ((Figure 3E) and (Benz et al., 2019). In the screen, the adapter protein non-catalytic region of tyrosine kinase (Nck) 2 bound selectively to the tyrosine phosphorylated VASP protein (Figure 3F). Nck2 is composed of three SH3 domains and one SH2 domain. Given that the proline-rich region of VASP directly interacts with SH3 domains (Benz et al., 2008; Benz et al., 2016), the SH3 domains of Nck2 could potentially mediate VASP-binding. Given, however that Nck2 only bound to phospho-tyrosine VASP, it is tempting to speculate that the SH2 domain of Nck2 directly interacts with phosphorylated tyrosine 16/39 in the VASP EVH1 domain. Interestingly, several studies highlighted the importance of Nck in angiogenesis (Dubrac et al., 2016; Lamalice et al., 2006) and Nck proteins are known to connect receptor (and non-receptor) tyrosine kinases to the machinery of actin reorganization, thereby regulating signal transduction of VEGF- and Eph-receptors and others (Alfaidi et al., 2021; Kania and Klein, 2016; Lettau et al., 2009). Thus, we speculate that complex formation of Nck2 and Ena/VASP proteins downstream of Eph receptors may participate in repulsion of endothelial tip cells or axonal growth cones (Figure 3G) through modulation of actin-rich protrusions, including lamellipodia and filopodia. Interestingly, Nck and VASP have been identified in a macromolecular complex that links the actin cytoskeleton to Fcγ receptor signaling during phagocytosis (Coppolino et al., 2001), suggesting that Nck-VASP binding may not be limited to signaling events downstream of Eph-receptors. Given that phagocytosis is a form of endocytosis, it is tempting to speculate that Nck-Ena/VASP complex formation in endothelial cells may also be implicated in membrane trafficking downstream of ephrin-EphB complex formation, which is critical to initiate cell repulsion (Pitulescu and Adams, 2010). Consistently, Ena/VASP-dependent receptor internalization/trafficking was demonstrated in the context of chemotaxis and guidance cue mediated cell migration (Laban et al., 2018; Tu et al., 2015; Vehlow et al., 2013; Zink et al., 2021) and a dominant negative approach to block Ena/VASP protein function impaired the internalization of phosphorylated EphB4 receptors in engineered fibroblasts (Evans et al., 2007).

**Figure 3.**
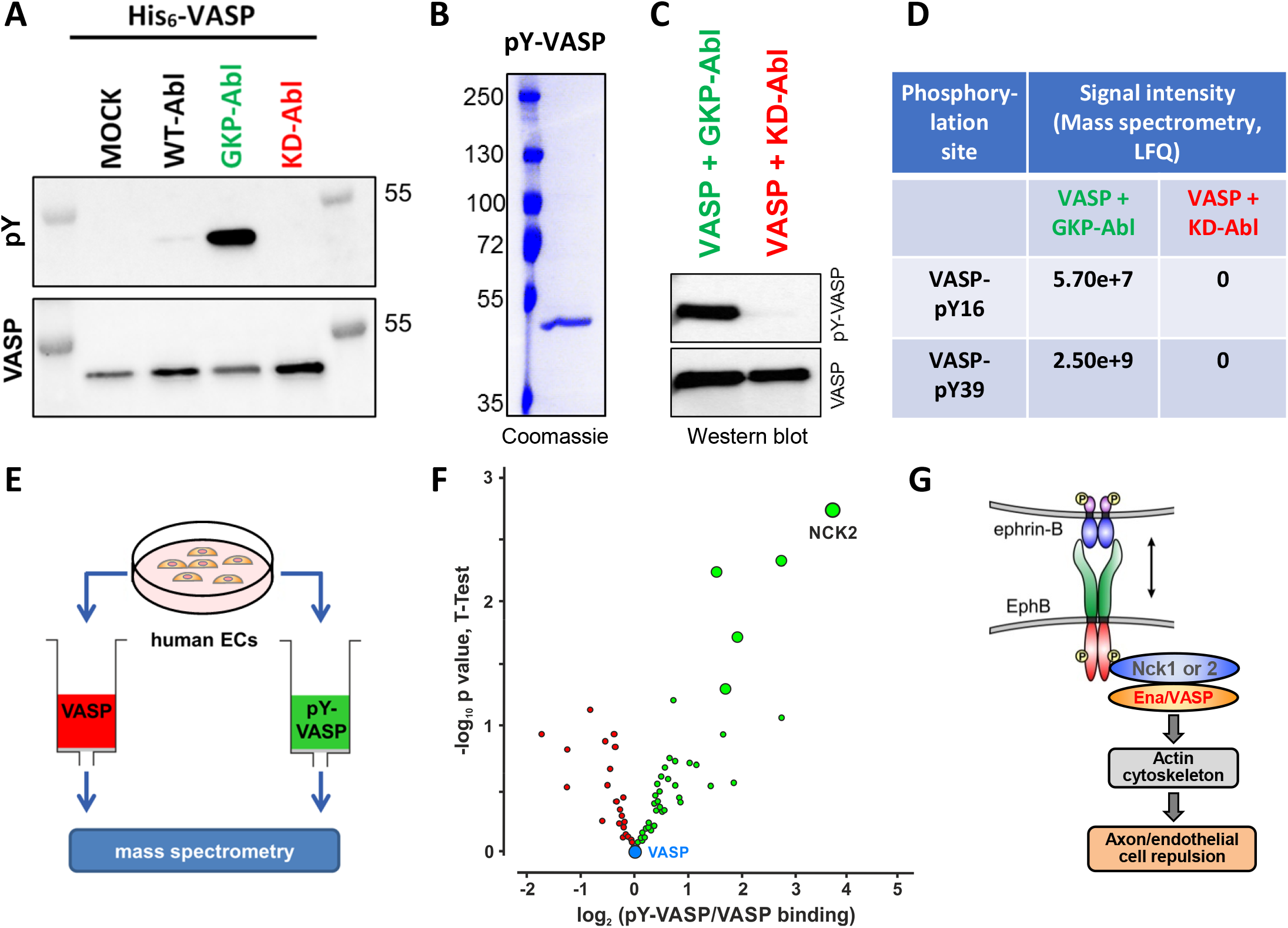
Identification of Nck2 as a novel phospho tyrosine-VASP binding protein. (**A**) HEK293 cells were co-transfected with His_6_-VASP and Abi-1, and empty vector (MOCK), WT-Abl, active GKP-Abl or inactive KD-Abl, respectively. Cell lysates were probed with phosphor-tyrosine specific antibodies (pY, 4G10, upper panel) or total VASP (lower panel). (**B**) Coomassie stained gel showing two micrograms of purified VASP from HEK293 cells co-transfected with GKP-Abl. Tyrosine phosphorylation status of purified VASP from HEK293 cells co-transfected with GKP-Abl or KD-Abl, respectively, was analyzed by Western blotting with phospho-tyrosine antibodies (pY, 4G10) (**C**) or by mass spectrometry (**D**); LFQ, label-free quantification. (**E**) Lysates of human endothelial cells (ECs) were incubated with immobilized VASP or tyrosine-phosphorylated VASP and differentially bound proteins were identified by mass spectrometry. (**F**) Volcano blot showing proteins from endothelial cell lysates that bound differentially to tyrosine-phosphorylated or non-phosphorylated VASP as identified by mass spectrometry. Proteins with highest fold change and statistical significance (p<0.05) are highlighted. The adapter protein non-catalytic region of tyrosine kinase (Nck) 2 bound selectively to the tyrosine-phosphorylated VASP protein. (**G**) Proposed model how Nck2 could link ephrin/EphB signaling to Ena/VASP-mediated actin remodeling and endothelial/axon repulsion.

## Materials and methods

Stripe assays were performed essentially as described (Knoll et al., 2007). Ephrin-B2 coating solution: human ephrin-B2-Fc (10 µg/ml, RnD Systems, #7397-EB) was mixed with Cy3-conjucated Fc-antibody (5 µg/ml, Sigma-Aldrich, # C2571) in HBSS and incubated for 30-60 minutes at room temperature with gentle agitation. Control stripe coating solution: IgG Fc-Fragment (10 µg/ml, Jackson ImmunoResearch, #009-000-008) was mixed with unconjugated Fc-antibody (5 µg/ml, Sigma-Aldrich, # I2136) in HBSS and incubated for 30-60 minutes at room temperature with gentle agitation. Stripe matrices were obtained from the Karlsruher Institut für Technologie (KIT; matrix code 2B, parallel 50 µm stripes). Solutions were subsequently applied to a glass slide for 45-60 minutes in a cell culture incubator separated by a washing step with HBSS. After coating of the control solution, the coated surface was incubated with fibronectin. Human umbilical vein endothelial cells (HUVECs) were isolated and cultured as described (Busse and Lamontagne, 1991). Cells were transfected with Mena and VASP siRNA (Qiagen SI00051373, SI02664200, SI 04208981, SI04366754) or control siRNA (Qiagen, #SI03650318) at 90% confluency (24 pmol siRNA per 12 well dish cavity with Lipofectamine RNAiMAX, Thermo Fisher Scientific). Sixty-five hours later, cells were detached, seeded on the ephrin-B2/control coated stripes and imaged with a Zeiss Cell Observer imaging system equipped with a 10x objective for 5 hours at 37°C and 5% CO_2_. One image every 5 minutes was recorded and compiled in one video. Cell movement in the videos was analyzed with ImageJ software using signal standard deviation as a measure of cell movement.

To induce EphB-stimulated tyrosine phosphorylation of VASP, HUVECs and HUAECs were starved for 3 hours in MCDB131 medium with 0.1% BSA for 3 hours. K562 cells were starved in RPMI 1640 medium without supplements for 3 hours. Cells were stimulated with clustered ephrin-B2-Fc or clustered Fc control for 30 minutes and then lysed in 1x TBS with 1% triton X-100 supplemented with protease and phosphatase inhibitors for 10 minutes on ice. After centrifugation, lysates were immunoprecipitated with IE273 anti-VASP antibodies (Immunoglobe) and protein G Sepharose (GE-Healthcare) for 1 hour at 4°C. After extensive washing in lysis buffer, precipitated material was eluted in SDS loading buffer and subjected to Western blotting with pY-specific antibody mix (pY20, pTyr100, 4G10) followed by secondary Mouse TrueBlot Ultra HRP anti-mouse IgG. Subsequently, membranes were re-probed with rabbit polyclonal anti-VASP antibodies (Benz et al., 2008). Protein loading was visualized by Ponceau-S staining.

His_6_-tagged human VASP was expressed and purified as described (Benz et al., 2009). Purified VASP was *in vitro* phosphorylated with active EphB2 (Sigma-Aldrich #E7157) in 20 mM Hepes-NaOH (pH 7.4), 5 mM MgCl2, 2.5 mM MnCl2, 1 mM EGTA, 0.4 mM EDTA, 0.05 mM DTT for 0-30 minutes at 30°C. GST-Zyxin (aa 1-143; in pGEX-4T3 vector) was expressed and purified as described (Benz et al., 2008) and covalently coupled to Affigel 15 (Biorad). 800 ng *in vitro* phosphorylated or non-phoshorylated VASP was pulled-down with 5 µg GST-Zyxin in a buffer containing 40 mM Hepes-NaOH pH 7.4, 75 mM NaCl, 1% Igepal CA-630 supplemented with protease and phosphatase inhibitors for 1 hour at 4°C. After extensive washing, precipitated material was analyzed by Western blotting with pY- or VASP-specific antibodies as detailed above.

Differential proteomics to identify pY-VASP/VASP binding proteins were performed as described (Benz et al., 2019).

## Supporting information

Supplementary movie 1_CTRL siRNA

Supplementary movie 2_Mena_VASP siRNA

## Acknowledgements

This work was supported by the Deutsche Forschungsgemeinschaft (SFB 834/A8 to PMB, SFB 834/A5 to IF). PMB was also supported by the German Center for Cardiovascular Research (DZHK B14-028 SE). The authors are indebted to Amparo Acker-Palmer for scientific input and help with the stripe assay.

